# Maximizing photon utilization in spectroscopic single-molecule localization microscopy using symmetrically dispersed dual-wedge prisms

**DOI:** 10.1101/2024.05.12.593746

**Authors:** Wei-Hong Yeo, Benjamin Brenner, Youngseop Lee, Junghun Kweon, Cheng Sun, Hao F. Zhang

## Abstract

Single-molecule localization microscopy (SMLM) enables super-resolution imaging on conventional fluorescent microscopes. Spectroscopic SMLM (sSMLM) further allows highly multiplexed super-resolution imaging. We report an easy-to-implement symmetrically dispersed dual-wedge prism (SDDWP)-sSMLM design that maximizes photon utilization. We first symmetrically dispersed photons to the −1^st^ and +1^st^ orders in an optical assembly using two identical dual-wedge prisms (DWPs). Then we computationally extracted the fluorophores’ spatial position and spectral characteristics using photons in both the −1^st^ and +1^st^ orders. Theoretical analysis and experimental validation showed lateral and spectral precisions of 10.1 nm and 0.3 nm, respectively, representing improvements of 28% and 48% over our previous DWP-based system, where emitted photons are divided separately for spatial and spectral analyses.

## 1. Introduction

Single-molecule localization microscopy (SMLM) offers a cost-effective solution over other competing super-resolution microscopy technologies like structured illumination microscopy (SIM) or stimulated emission depletion (STED), as it mainly relies on conventional fluorescence microscopes coupled with specialized software for image acquisition and analyses [1]. The cost-effectiveness of SMLM has not only increased its accessibility to researchers across diverse fields but has also spurred community interest in advancing the technology further. Driven by the demand for multiplexed imaging, several modifications to SMLM have been developed. These include employing dichroic filters to segregate different fluorophores by their spectral band [2] or utilizing ratiometric and unmixing methods based on intensity differences between the two detection channels [3]. However, these methods have restricted the number of color channels that can be simultaneously imaged. Alternatively, increased multiplexing can be achieved by sequentially activating different fluorophores at the expense of prolonged acquisition times with repeated sample unlabeling and relabeling [4, 5].

Spectroscopic SMLM (sSMLM) overcomes these limitations by introducing a dispersive element, such as a diffraction grating [6, 7] or a prism [8, 9], to simultaneously image the spatial position and spectral characteristics of individual fluorophores. By capturing and analyzing the complete emission spectrum, sSMLM has theoretically unlimited flexibility in the number and combination of fluorophores for high throughput parallel multiplexed imaging. Operating as a shot-noise limited system, its precision in localization and spectral identification relies on effectively utilizing the limited number of photons emitted in each single molecular emission process [10]. Earlier sSMLM utilized diffraction gratings to separate photons into the 0^th^ and 1^st^ order diffractions for spatial localization and spectral identification [11]. However, allocating photons to spectral channels reduced the localization precision in sSMLM.

To improve the efficiency of photon utilization in sSMLM, we previously introduced the symmetrically dispersed sSMLM (SDsSMLM) design. Although this method uses a diffraction grating, it leverages the mirror symmetry of the two spectral channels (−1^st^ and +1^st^ orders) to effectively use the photons for both spatial and spectral analysis, improving spatial and spectral uncertainties by 42% and 10%, respectively, compared to implementations using the 0^th^ and 1^st^ orders [12]. To overcome the issue of low transmission efficiency in gratings, we further developed a dual-wedge prism (DWP)-based sSMLM. This approach utilizes a monolithic assembly of a beamsplitter and a DWP, which reduces aberrations and photon loss more effectively than a diffraction grating. As a result, it enhanced lateral precision by 47% and axial precision by 23%, while maintaining a comparable spectral precision [13].

The combined benefits of these two technologies promise an ideal solution for maximizing photon utilization, thereby further enhancing the performance of sSMLM. Here, we report symmetrically dispersed dual-wedge prism sSMLM (SDDWP-sSMLM). SDDWP-sSMLM employs a pair of DWPs, effectively dispersing fluorescent emissions into both the −1^st^ and +1^st^ spectral channels. This ensures full utilization of collected photons for both spatial localization and spectral identification. Notably, the device retains its monolithic form. It is easily attachable to the side port of an inverted optical microscope, thus converting a conventional fluorescent microscope into sSMLM without needing additional optical alignment or adjustment steps. This cost-effective plug-in optical assembly module simplifies dissemination to a broader research community.

In this paper, we first construct analytical models to simulate the characteristic performance of SDDWP-sSMLM for two-dimensional (2D) imaging. We then experimentally measure the lateral and spectral precision using a nano-fabricated calibration target composed of a 2D array of nanoholes machined into an aluminum film on a glass substrate illuminated by a tunable laser source. Upon confirming the accuracy of the analytical model through experimental data, we extend our investigation to include axial offsets of two spectral channels for three-dimensional (3D) imaging, updating the analytical models accordingly. Finally, we demonstrate the practicality of imaging single cells using 3D SDDWP-sSMLM.

## 2. Methods

### 2.1. Sample Preparation

We used a custom-made nanohole array for a static calibration target for spectral calibration. First, a 150 nm thick aluminum (Al) film was sputtered onto a borosilicate glass coverslip measuring 22 mm × 22 mm (ATC Orion Sputter System; AJA International) with a direct current power of 75 W and an argon flow rate of 20 standard cubic centimeters per minute (sccm) at 3 mTorr. Subsequently, 5 by 8 Al nanohole arrays, featuring a rectangular lattice with a period of 4 µm and a hole diameter of 100 nm, were created using focused ion beam (FIB) milling (JIB-4700F FIB-SEM; JEOL Ltd.). Figure S1 show an SEM image of the nanohole array.

For axial calibration, we prepared fluorescent microsphere samples (F8807; ThermoFisher) diluted approximately 10^6^ times in PBS to give us single microsphere samples. We then deposited the microspheres on an 8-well chambered coverglass (12-565-470, Fisher Scientific) to match the imaging conditions of the cellular samples.

To validate our experimental setup using biological samples, we used a multiplexed labeling approach to visualize specific intracellular structures within HeLa cells. Specifically, we used DY-634, Alexa Fluor 647 (AF647), and CF660C dyes to target peroxisomes, microtubules, and mitochondria, respectively. We permeabilize the cells for 10 minutes using a solution containing 0.2% Triton X (Sigma Aldrich; X100) in phosphate-buffered saline (PBS) (ThermoScientific; J61196). Subsequently, a blocking step was performed, which involved a 30-minute incubation in a blocking buffer composed of 2.5% Normal Goat Serum (NGS) (ThermoFisher; 50062Z) and 2.5% Bovine Serum Albumin (BSA) (Fisher Bioreagents; 9048-46-8) in PBS.

The primary antibody incubation was carried out at 4 ^°^C overnight, where we used a mix of antibodies including Mouse Anti-PMP70 (Sigma-Aldrich; SAB4200181; 1:300 dilution), Rabbit Anti-Tom20 (MilliporeSigma; HPA011562; 1:200 dilution), and Rat Anti-beta-Tubulin (MilliporeSigma; MAB1864; 1:150 dilution). All the antibodies were diluted in the same blocking buffer. After primary antibody incubation, we thoroughly washed the cells thrice for 5 minutes each using a washing buffer solution of 2% NGS, 2% BSA, and 0.1% Triton X in PBS.

Subsequently, we incubated conjugated secondary antibodies targeting the primary antibodies in the blocking buffer at a 1:200 dilution for 45 minutes at room temperature. Conjugated secondary antibodies (JacksonImmunoResearch; Anti-Sheep: 713-005-147, Anti-Mouse: 715-005-151, Anti-Rat: 712-005-153) were made by reacting NHS-ester dyes with the antibodies, and the dyes used were DY-634 (Dyomics; 634-01), AF647 (ThermoFisher; A20006), and CF660C (Biotium; 92137), respectively. After this incubation, we washed the cells again twice for 5 minutes each using the washing buffer and then twice for 5 minutes each using PBS.

### 2.2. SDDWP-sSMLM imaging setup

The SDDWP-sSMLM imaging setup was built on our previously reported sSMLM system [13], which uses an inverted microscope body (Ti2-E; Nikon) with a 100× total-internal reflection fluorescence (TIRF) objective (MRD01995; Nikon). Figure 1(a) illustrates the light path of our microscope setup. For fluorescent samples, we used a 642 nm excitation laser (2RU-VFL-P-2000-642-B1R; MPB Communications), which passes through a bandpass filter (BPF, FF01-640/14-25; Semrock) and then reflected by a dichroic mirror (DM, Di03-R635-t1-25x36; Semrock) before illuminating the sample. The fluorescent sample then fluoresces at a red-shifted wavelength, where it is then captured by the objective, passes through the DM, and is filtered by a long-pass filter (LPF, BLP01-635R-25; Semrock). After passing through a focusing tube lens, the detected fluorescence is then passed to either the imaging configuration of 2D (Fig. 1(b)) or 3D (Fig. 1(c)) SDDWP-sSMLM.

**Fig. 1.**
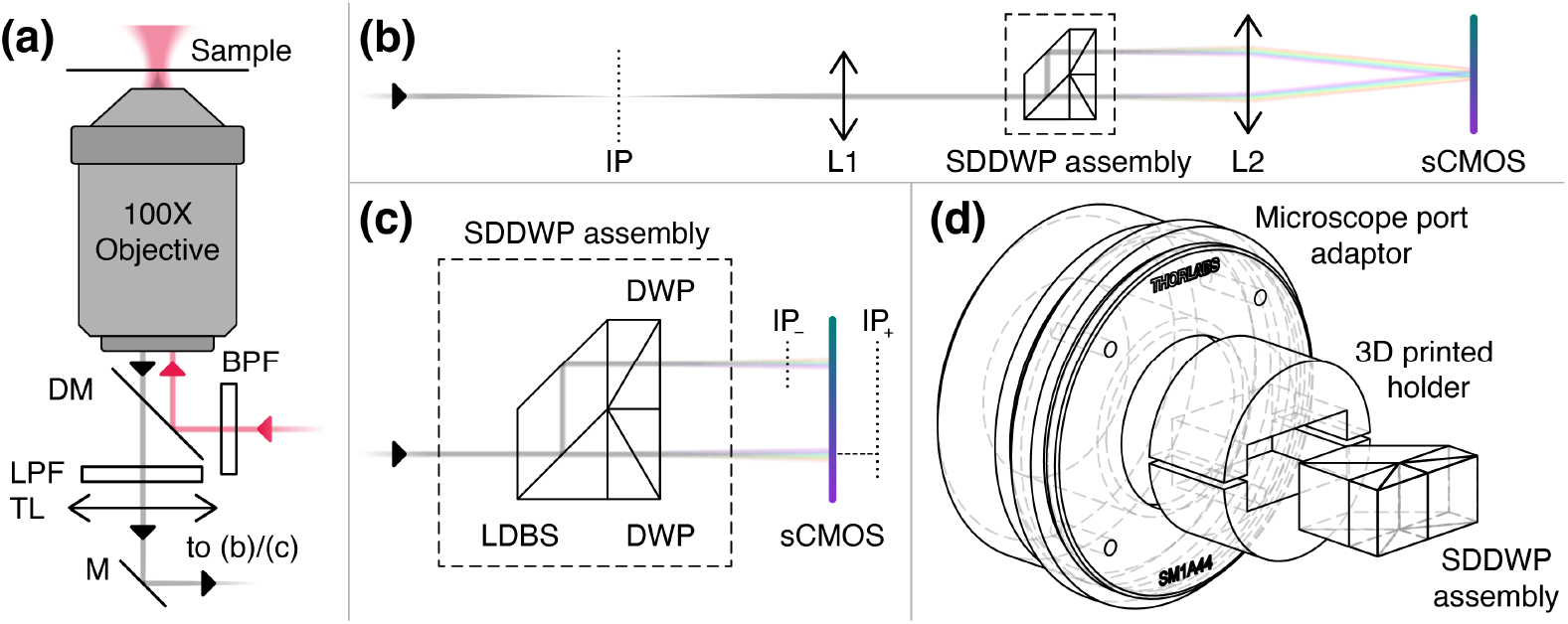
SDDWP-sSMLM working principle. (a) Beam paths of the excitation laser used for imaging samples (red) and the emitted photons from the sample (gray); (b) 2D SDDWP-sSMLM detection setup; (c) 3D SDDWP-sSMLM detection setup. The dotted box is a blown-up view of the SDDWP assembly; (d) Illustration of the microscope port adaptor, 3D printed holder, and SDDWP assembly.

For 2D SDDWP-sSMLM shown in Fig. 1(b), we used a matching pair of L1 and L2 (*f* = 100 mm, 49-390; Edmund Optics) to form a 4F optical relay. It projects the imaging plane (IP) onto the scientific complementary metal-oxide semiconductor (sCMOS) sensor (Prime 95B 25mm; Photometrics), while the SDDWP assembly is located near its common focal plane. Figure 1(c) shows a magnified illustration of the SDDWP assembly, where the lateral displacement beamsplitter (LDBS) splits the image into two, and in each beam path, we incorporated a DWP to provide the necessary spectral dispersion for detection. The DWP specifications used here follow our previous publication [13] (Hyperion Optics). For 3D SDDWP-sSMLM, the SDDWP assembly splits the image into two, then focuses each image from the microscope onto image planes IP_−_ and IP_+_ due to the difference in beam paths shown in Fig. 1(c), forming a typical biplane setup [14, 15]. An sCMOS camera simultaneously captured photons from both beam paths for further processing. Finally, the holder for the SDDWP assembly was custom-designed, 3D printed, and mounted onto a microscope camera port adapter (SM1N2; Thorlabs), as shown in Fig. 1(d).

For the non-fluorescent nanohole array used in lateral and spectral calibration, we used the fluorescent LED overhead (TI2-D-LHLED; Nikon) included with our Ti2-E microscope to illuminate the sample and removed the DM and LPF along the imaging path. We then added a filter wheel that holds five bandpass filters with center wavelengths of 532 nm, 580 nm, 633 nm, 680 nm, 750 nm (FL532-3, FBH580-10, FLH633-5, FBH680-10, and FBH750-10; Thorlabs) to select the desired bandwidth of light for calibration. For quantifying the localization and spectral precisions, we used a supercontinuum (SC) laser (SC-PRO 7; YSL) with an acousto-optic tunable filter (AOTF) (NKT Photonics) with the light source placed above the sample to adjust the illumination wavelength precisely. Again, we removed the DM and LPF along the imaging path and imaged them without the filter wheel.

### 2.3. SDDWP-sSMLM calibration and imaging protocols

#### 2.3.1. Calibration protocol

We first used our nanohole array to calibrate the relative lateral positions in both the −1^st^ and +1^st^ orders image against the known hole positions verified by SEM. This corrects for magnification errors present due to pathlength differences. Here, we used the LED lamp to illuminate the nanohole array, and added the 680 nm filter in the emission path to select this bandwidth of light, as it closely matches the emission wavelengths used in our cell sample. We captured 100 frames on the sCMOS with an exposure time of 100 ms.

To calibrate the positions of localizations after spectral dispersion relative to different known wavelengths, we captured images of the nanohole array through the five bandpass filters with the sCMOS for 100 frames with exposure times adjusted to provide sufficient signal-to-noise ratios. Lastly, we imaged fluorescent microspheres at different *z*-positions for axial calibration, ranging from −2.0 µm to 2.0 µm in 20 nm steps. We used the 642 nm excitation laser to excite the microspheres and captured the 201 frames using the sCMOS camera.

#### 2.3.2. Imaging protocols

We quantified our system’s lateral and spectral precision by illuminating the nanohole sample with the SC laser and AOTF set to 625 nm and used the sCMOS camera to capture 1,000 frames with an exposure time of 10 ms per frame. As a static sample, the nanohole array also facilitated drift corrections for our spatial precision measurements.

For experimental validation, we imaged our cell samples prepared with the protocol outlined in Section 2.1 at 642 nm excitation with a power density of ∼10 mW/mm^2^. We used the sCMOS camera to capture 30,000 frames with an exposure time of 10 ms per frame. We imaged the same sample at 3 different *z*-positions with offsets of −0.3 µm, 0.0 µm, and +0.3 µm to give a better axial visualization of the cell sample.

### 2.4. SDDWP-sSMLM image processing

To localize the fluorophores or the spots of the nanohole array, we used the ThunderSTORM ImageJ plug-in [16], providing us with a localization table. We localized the −1^st^ and +1^st^ orders separately, giving us localization information of the *x*- and *y*-coordinates, including other useful parameters such as intensity and width of the localizations. Custom data processing code for manipulating and matching the localizations was written in MATLAB R2021a. The lateral, axial, and spectral information was inferred from the localizations using the methods described in Section 2.4.1.

#### 2.4.1. Inferring lateral, axial, and spectral information

As illustrated in Fig. 2(a), after localizing the fluorophores, we obtain key parameters *x*_±_, *y*_±_, *σ*_*x*, ±_ and *σ*_*y*, ±_ from the real ±1^st^ order images. See Table 1 for nomenclature definitions. First, we set *x*_0_ and *y*_0_ as the ‘true’ localizations of the *x*- and *y*-coordinates, respectively, where they form a virtual image shown in the middle of the real −1^st^ and +1^st^ order images. We can define general relations for the localizations of the −1^st^ and +1^st^ order images:

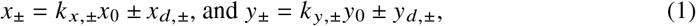

where *k* _*x*,±_ and *k* _*y*,±_ are the magnification factors in the *x*- and *y*-directions, respectively, and *x*_*d*,±_ and *y*_*d*,±_ are the lateral shifts induced by the SDDWP assembly along the *x*- and *y*-directions for the −1^st^ and +1^st^ orders. To simplify the analysis, we make three assumptions: (1) our magnification factors are equal and unity, i.e. *k* _*x*,±_ = *k* _*y*,±_ = 1, (2) the SDDWP assembly has perfect symmetry along the *x*-direction, i.e. *x*_*d*,−_ = *x*_*d*,+_ = *x*_*d*_, and (3) there are no lateral shifts along the *y*-direction, i.e. *y*_*d*,−_ = *y*_*d*,+_ = 0, since we orientate the SDDWP assembly to maximize dispersion along the *x*-direction. Thus, the equations simplify to *x*_±_ = *x*_0_± *x*_*d*_ and *y*_±_ = *y*_0_. To obtain the lateral information from the −1^st^ and +1^st^ order images, we take the linear combinations of Eq. (1), which gives us *x*_0_ = (*x*_−_+*x*_+_)/2, and *y*_0_ = (*y*_−_+*y*_+_)/2, respectively, which is simply the average between the two localizations, as *x*_*d*_ and *y*_*d*_ cancel out due to symmetry.

**Table 1.** Nomenclature used in this paper.

| Notation | Description |
| --- | --- |
| $x, y, z$ | $x$-, $y$-, $z$-coordinates of a given localization |
| $x_0, y_0$ | $x$-, $y$-coordinates of the reconstructed localizations |
| $x_{\pm}, y_{\pm}$ | $x$-, $y$-coordinates of the localizations of the $-1^{\text{st}}$ and $+1^{\text{st}}$ order images |
| $k_{x,\pm}, k_{y,\pm}$ | Magnification factors of the $-1^{\text{st}}$ and $+1^{\text{st}}$ order images in the $x$- and $y$-directions |
| $x_{d,\pm}, y_{d,\pm}$ | Lateral dispersion of the $-1^{\text{st}}$ and $+1^{\text{st}}$ order images in the $x$- and $y$-directions due to the SDDWP assembly |
| $\sigma_{x,\pm}, \sigma_{y,\pm}$ | Fitted gaussian widths of the $-1^{\text{st}}$ and $+1^{\text{st}}$ order images in the $x$- and $y$-directions |
| $x_{\lambda}$ | Dispersion due to a DWP in the $x$-direction |
| $\lambda$ | Wavelength of an emitter |
| $f(\lambda)$ | Invertible empirical function that corrects for the non-linearity of DWP dispersion |
| $m$ | Scaling constant for the dispersion |
| $c$ | Offset constant for the lateral shift in positions between the $-1^{\text{st}}$ and $+1^{\text{st}}$ order localizations |
| $l$ | Mutual defocus between the two channels provided by the difference in the beam path lengths due to the LDBS |
| $d$ | Scaling constant for the rate of change of defocus of the width of the PSF with $z$-position |
| $F$ | Ratio of defocus curves for axial calibration |
| $N$ | Number of emitted photons |
| $a$ | Pixel size of the imaging sensor |
| $b_{\text{bg}}, b_{\text{ro}}$ | Mean number of background photons per pixel and the standard deviation of readout noise |
| $\tau$ | Background correction factor |
| $\Delta x_{-}, \Delta x_{+}$ | Lateral localization precisions of the $-1^{\text{st}}$ and $+1^{\text{st}}$ order images |
| $\Delta x, \Delta z, \Delta \lambda$ | Lateral, axial and spectral precisions |

**Fig. 2.**
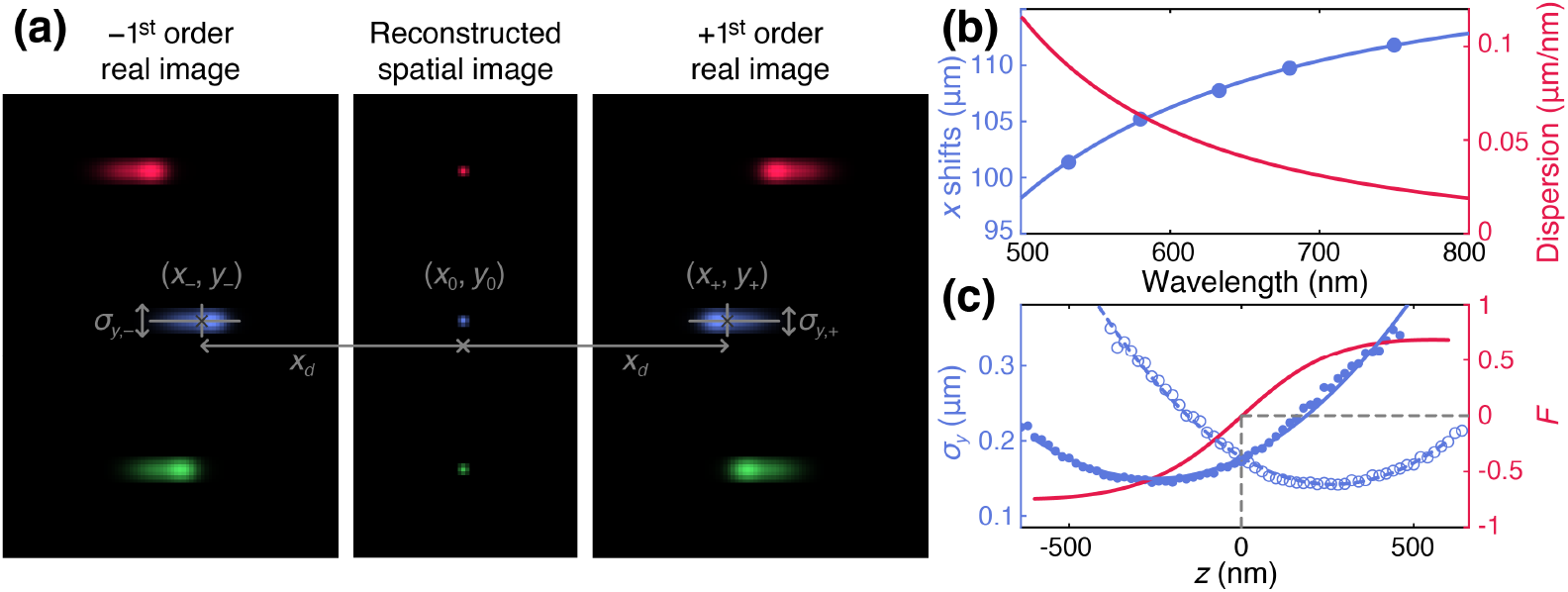
(a) Illustration of the resulting −1^st^ and +\1^st^order real images on the sCMOS, with the reconstructed virtual spatial image in the middle; (b) The blue line shows the spectral calibration curve of the DWP assembly, which is used to determine the characteristic wavelength of each localization. The red line shows the dispersion of the system, which is the instantaneous gradient of the blue curve; (c) The axial calibration curve used to determine the axial location of each localization. The blue lines illustrate the width of different focal spots associated with each *z*-position set on the microscope, and the red S-curve is used to infer the *z*-position of the localizations.

Since the dispersion resulting from the DWP is non-linear, we first use a rational invertible empirical function, *f* (*λ*) [17], to correct its non-linearity and provide a simple linear relation between observed dispersions with a DWP, *x*_*λ*_, and *f* (*λ*):

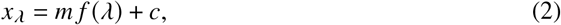

where *m* is a measure of spectral dispersion of a DWP, and *c* accounts for any lateral shift. We can apply Eq. (2) to both the −1^st^ and +1^st^ order images to obtain *x*_*d*,±_ = *m*_±_ *f* (*λ*) + *c*_±_. Now, to determine the characteristic wavelength of each localization, we compute *x*_+_ − *x*_−_, which is the difference between the lateral positions of the −1^st^ and +1^st^ order images, to get

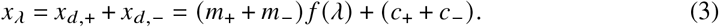

First, Eq. (3) retains the same form as Eq. (2), implying linearity. Since we assumed that *x*_*d*, −_ = *x*_*d*,+_ = *x*_*d*_ above, it follows that the spectral calibration parameters must be equal, i.e. *m*_+_ = *m*_−_ and *c*_+_ = *c*_−_. If we set *m* = *m*_+_+ *m*− and *c* = *c*_+_ +*c*_−_, we may determine the constants *m* and *c* with a spectral calibration procedure described in Section 2.3.1. We can then determine the characteristic spectra of a given emitter by inverting Eq. (2) to get

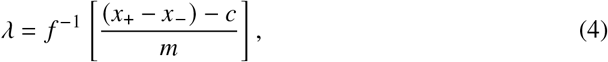

where *f* ^−1^ is the inverse of the invertible function *f*.

For 3D SDDWP-sSMLM, the biplane method is used to obtain axial information, as previously reported [13–15, 18]. A defocus curve was used to relate the axial position of the emitter with the spot size, *σ*_*y*, ±_, and the mutual defocus, 2*l*, between the two channels provided by the difference in the optical path lengths due to the LDBS. Specifically, the defocus curve is given by

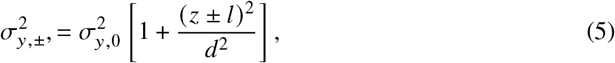

where *σ*_*y*,0_ is the spot size in focus, and *d* and *l* are constants to be determined through axial calibration (see Section 2.3.1). To obtain the axial position associated with the emitter, we have

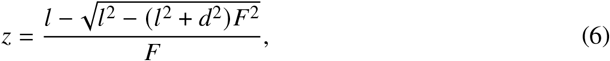

where 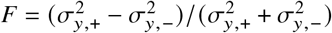.

#### 2.4.2. Lateral, spectral, and axial calibration

For lateral calibration, we matched the −1^st^ and +1^st^ order images to our known nanohole spacing intervals. This allows us to have an accurate measure of the lateral positions when determining *x*_0_ and *y*_0_ from Eq. (1). To perform spectral calibration, we compared lateral displacements of the nanohole array with different bandpass filters inserted, each cutting off the illumination of the LED lamp with a specific peak wavelength. By performing a best-fit to Eq. (2) (Fig. S2), we identified the constants *m* and *c* used to create the spectral calibration graph shown in Fig. 2(b). For axial calibration, we used the widths of the localizations along the *y*-direction in the −1^st^ and +1^st^ order images and performed a best-fit of Eq. (5), setting the axial position using the microscope. This provided us with the values of *d* and *l* needed for evaluating the relative axial position in Eq. (6), as shown in Fig. 2(c).

#### 2.4.3. Quantification of lateral and spectral precisions

To assess SDDWP-sSMLM’s performance, we quantified our system’s lateral and spectral precisions. First, we determined the lateral localized *x*- and *y*-positions of the −1^st^ and +1^st^ order images after correcting for sample drift. To correct for drifts in the nanohole sample, we computed the average position of all localized emitters in each frame to get a sample drift curve. We then subtracted the localized positions from the sample drift curve to obtain a drift-corrected localization table, which was then used to compute lateral precision by taking the standard deviation of the result from Eq. (1). For the spectral precision, we computed the standard deviation of the calculated spectral peak from Eq. (4) for each emitter.

#### 2.4.4. Multi-color 3D imaging processing

To demonstrate the capability of cellular imaging, we imaged HeLa cells prepared as described in Section 2.1 and imaged them as described in Section 2.3.2. For each of the three movies captured, we processed the frames using ThunderSTORM to obtain the localizations of the fluorophores in the −1^st^ and +1^st^ orders. We then used a custom routine written in MATLAB R2021a to read the localizations from ThunderSTORM and match the localizations between both orders. This script uses the axial calibration curve to determine the axial position of the fluorophores from the *y*-widths of the localizations and the spectral calibration curve to determine the spectral identity from the relative positions of the fluorophores. We then exported this file and performed drift corrections in ThunderSTORM to give us a fully processed, drift-corrected image for each *z*-offset. Finally, to reconstruct the image, we used another custom script that aligns each *z*-offset’s localizations and applies the offset to produce a combined localization output file.

## 3. Results and Discussion

### 3.1. Validation of 2D SDDWP-sSMLM

We evaluated the improvement provided by SDDWP-sSMLM over our previous DWP-sSMLM by comparing the lateral and spectral precisions. Such a comparison is best performed using a 2D SDDWP-sSMLM, where both imaging orders are in focus to determine the theoretical maximum precision attainable with SDDWP-sSMLM.

#### 3.1.1. Lateral precision

The primary factor limiting the lateral localization precision is the number of emitted photons, *N*, as fluorophores usually have a limited photon budget before photobleaching [10]. Thus, maximizing photon utilization improves spatial localization precision and image quality. The spatial lateral localization precision, Δ*x*, of a single localization is determined by [18]

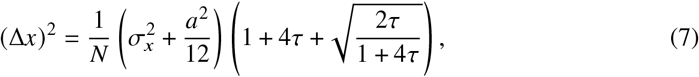

where *a* is the pixel size, *σ*_*x*_ is the standard deviation of the point spread function (PSF) of the spot, and *τ* is the background correction factor defined as 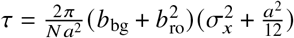, where *b*_bg_ and *b*_ro_ are the mean number of background photons per pixel and the standard deviation of readout noise, respectively. We further define Δ*x*_−_ as the lateral localization precision for the 0^th^ and −1^st^ order image for DWP- and SDDWP-sSMLM, respectively, and Δ*x*_+_ to denote the localization precision of the +1^st^ order image for both DWP- and SDDWP-sSMLM.

In DWP-sSMLM, we used a DWP assembly to generate a 0^th^ order image for lateral localization and a +1^st^ order image for spectral determination. Since only the 0^th^ order image is used for localization in DWP-sSMLM, only 45% of the overall photon budget is available for lateral localization. In comparison, SDDWP-sSMLM doubles the overall photons used for lateral localization, because it uses photons captured in both the −1^st^ and +1^st^ order images. Therefore, the effective lateral precision of an emitter in SDDWP-sSMLM is defined as

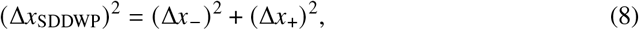

where Δ*x*_−_ and Δ*x*_+_ are the lateral localization precisions of the −1^st^ and +1^st^ order images of SDDWP-sSMLM, respectively. Since the photon budget *N* is approximately equal in both the −1^st^ and +1^st^ order images, we have Δ*x*_−_ ≈ Δ*x*_+_, and thus 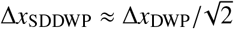.

The precision improvement is shown in Fig. 3(a), where the lateral precision of SDDWP-sSMLM (solid red curve) is 27% better than that of DWP-sSMLM (solid blue curve). Such an improvement is also evident by our experimental measurements, as shown by the red and blue dots in Fig. 3(a), indicating good agreement between our experimental measurements and theoretical predictions, with an average error of 0.7 nm and 0.8 nm for DWP- and SDDWP-sSMLM, respectively. At a photon count of 2,000, the lateral precisions are 6.4 nm and 4.7 nm for DWP- and SDDWP-sSMLM, respectively, representing an improvement of 34%.

**Fig. 3.**
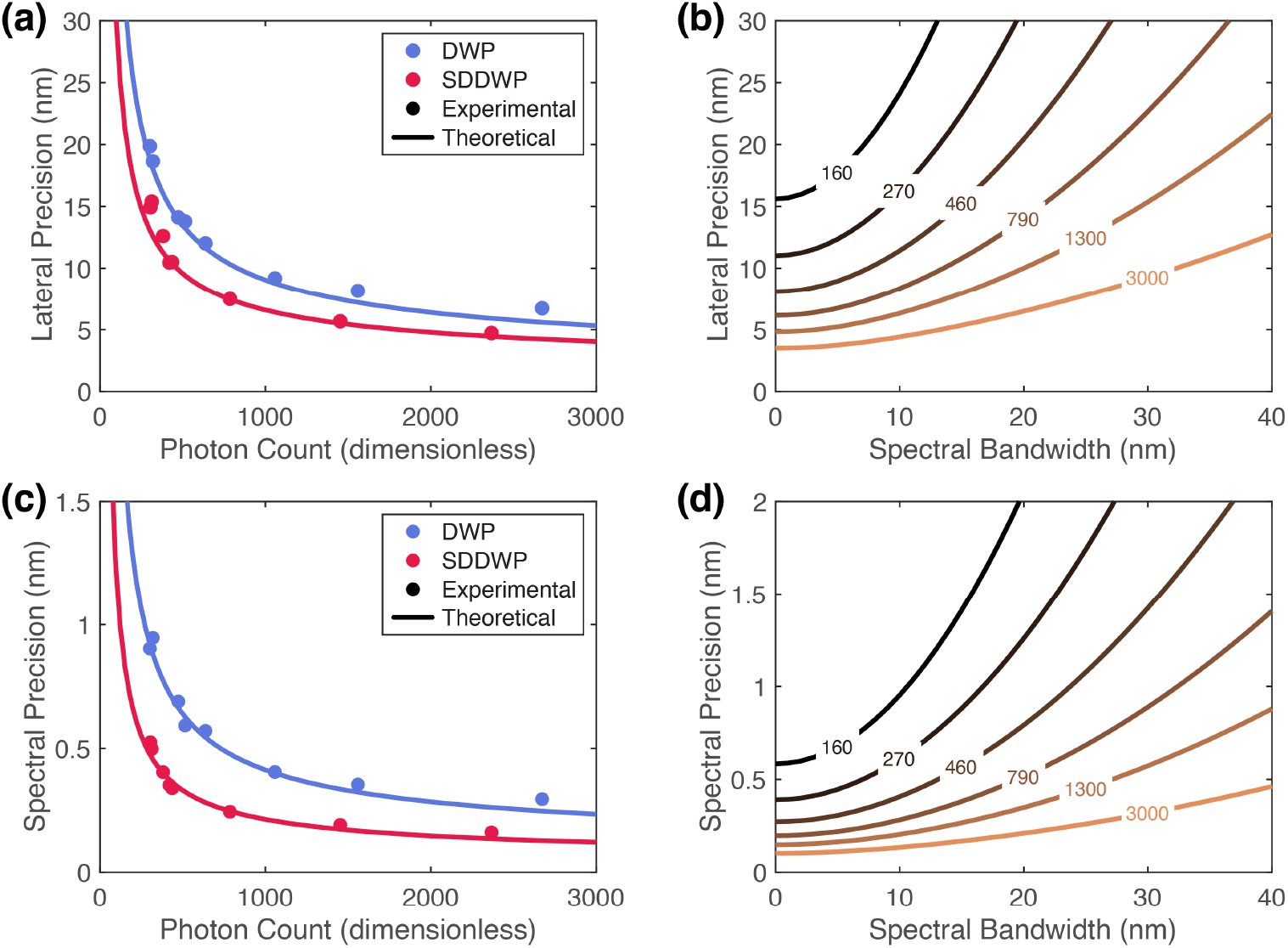
(a) Comparison of lateral precisions between SDDWP- and DWP-sSMLM; (b) Theoretical lateral precision with different emission spectral bandwidths and photon counts. The numbers along the lines represent the photon counts; (c) Comparison of spectral precision between SDDWP- and DWP-sSMLM; (d) Theoretical spectral precision for emission spectral bandwidths and photon counts.

Using the SC laser with the AOTF, the typical FWHM emission spectra is 5.6 nm. Therefore, the dispersion along the *x*-direction is limited, and the lateral localization precision is relatively unaffected according to Eq. (7). Indeed, the differences in the lateral localization precision between the *x*- and *y*-directions are 0.8 nm and 1.0 nm for DWP- and SDDWP-sSMLM, respectively. This is confirmed by Fig. 3(b), where a spectral bandwidth of 5 nm does not affect lateral precision significantly.

#### 3.1.2. Spectral precision

To obtain the analytical expression for spectral precision, we performed a Taylor expansion on Eq. (4) to obtain

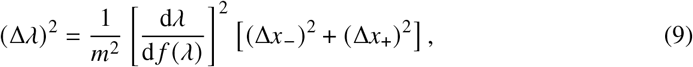

where 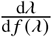 is the local spectral dispersion at the central wavelength, and *m*, Δ*x*_−_, and Δ*x*_+_ are the same as previously defined.

In SDDWP-sSMLM, we get twice the amount of spectral dispersion compared to DWP-sSMLM due to two DWPs. This can be seen in Eq. (3), where for DWP-sSMLM, we have *m*_−_ = 0 due to the absence of a DWP in the 0^th^ order image. Therefore, we get *m*_SDDWP_ = 2*m*_DWP_, and thus (Δ*λ*)_SDDWP_ = (Δ*λ*)_DWP_ /2. In Fig. 3(c), we plot the theoretical curve governed by Eq. (9) for DWP- and SDDWP-sSMLM. As discussed in Section 3.1.1, there is a spectral bandwidth of 5.6 nm in our SC laser and AOTF setup, which increases the value of *σ*_*x*_, and consequently, Δ*x*_−_ and Δ*x*_+_. From Eq. (7), we see that this gives us a higher value of spectral precision. Therefore, to account for this effect, we adjusted the uncertainties of Δ*x*_−_ and Δ*x*_+_ with the spectral bandwidth measured and the resulting graphs are shown in Fig. 3(c). The theoretical estimations agree well with the experimental results, with a mean errors of 0.03 nm and 0.02 nm for DWP- and SDDWP-sSMLM, respectively. Overall, we see a 34% improvement in spectral precision compared to DWP-sSMLM. At a photon count of 2,000, we see a 45% improvement in

##### SDDWP-over DWP-sSMLM

Similar to lateral precision, spectral precision is also affected by the spectral bandwidth of the emission. In Fig. 3(d), we plotted the theoretical spectral precisions for different spectral bandwidths, and we see that the spectral precision is higher under narrower spectral bandwidths. For most fluorophores used in sSMLM, the spectral bandwidth is typically less than 20 nm, and therefore, the spectral precision is generally better than 1 nm.

### 3.2. Non-ideal analyses

Ideally, the −1^st^ and +1^st^ order images are perfectly symmetrical; however, in practice, the two images may have slightly different magnifications, and the dispersions in the −1 ^st^ and +1^st^ order images are not identical. Here, we relax each assumption made in Section 2.4.1 and provide insights into the impact of these relaxations on the lateral and spectral calculations.

#### 3.2.1. Non-ideality in magnifications

Due to the pathlength difference between the −1^st^ and +1^st^ order images, there is a difference between the magnification factors of the two images. Accounting for this difference, we have

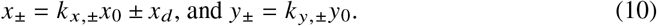

Thus, we may determine *x*_0_ and *y*_0_ by taking a weighted average of the −1^st^ and +1^st^ order images,

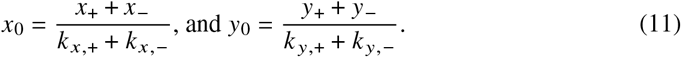

The resulting *x*_0_ and *y*_0_ are affected by the magnification differences by a scaling factor (*k*_*x*, +_ + *k* _*x*, −_) /2 and (*k* _*y*, +_ + *k* _*y*, −_) /2, respectively. We found from Fig. S3 that *k*_*x*,+_ / *k* _*x*,−_ = 1.012, and *k*_*y*, +_/*k* _*y*,−_ = 1.005. This gives us a lateral accuracy error of 0.6% and 0.3% for the *x*- and *y*-directions of reconstructed virtual images compared to uncorrected images of SDDWP-sSMLM.

Next, to compute the spectral information, we use the difference in the lateral positions of the −1^st^and +1^st^order images, *x*_*λ*_ = *x*_+_ −*x*_−_, which is now affected by the magnification differences. We have

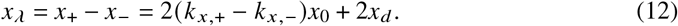

We observe that the error in *x*_*λ*_ is linear with respect to *x*_0_. This indicates a systematic error in the spectral calculations when the magnification difference between the −1^st^ and +1^st^ order images is not considered. For small fields of view, this error is negligible since the range of *x*_0_ is small, but for larger fields of view, this error becomes significant.

#### 3.2.2. Non-ideality in DWP dispersions

Due to differences in the dispersion of the −1^st^ and +1^st^ order images from manufacturing tolerances, the spectral calibration curve may not be the same for both images. Accounting for this difference, we have

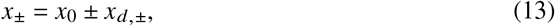

where *x*_*d*, −_ and *x*_*d*, +_are the dispersions of the −1^st^ and +1^st^ order images, respectively. From Eq. (3) and the discussion in Section 2.4.1, we can directly determine the spectral calibration parameters *m* and *c* for SDDWP-sSMLM, since Eq. (2) is linear.

Next to determine *x*_0_, we take the sum of the −1^st^ and +1^st^ order images, which gives us

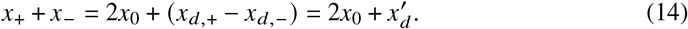

Observe that we have from Eq. (3) that 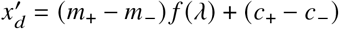, which is again in the form of Eq. (2). Rewriting in terms of *x*_0_, we have

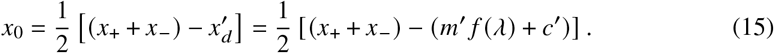

with *m*^′^ = *m*_+_−*m*_−_ and *c*^′^ = *c*_+_−*c*_−_. This implies that the error in *x*_0_ is linear with respect to the wavelength of the emitter. In other words, if the dispersion difference between the −1^st^ and +1^st^ order images is not considered, there is a systematic error in the spectral calculations. In Fig. S2, we show the fitted parameters of the −1^st^ and +1^st^ order images and the resulting spectral calibration curve for SDDWP-sSMLM. Although there is a slight difference in the dispersion of the −1^st^ and +1^st^ order images affecting lateral accuracy, this can be corrected for by determining 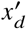 from a spectral calibration curve.

### 3.3. Performance analysis of 3D SDDWP-sSMLM

In 3D SDDWP-sSMLM, we combined the −1^st^ order and +1^st^ order images with biplane imaging for 3D localization. As a result, images from the two orders would be defocused differently, leading to enlarged PSFs, and thus higher Δ*x*_±_ in 3D than 2D imaging. Consequently, the lateral and spectral precisions are poorer than those of 2D imaging. To simplify our analysis, we set the photon count to 2,000, which is typical in sSMLM.

#### 3.3.1. Axial precision

With biplane imaging, we extend the capability of localizing fluorophores along the *z*-direction. First, we note that the biplane method is identical for both DWP- and SDDWP-sSMLM by estimating the width of the PSFs along only the *y*-direction, which is independent of the axis of spectral dispersion. To obtain the expression for axial precision, we start with [18]

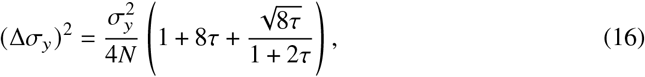

where *τ* is the background correction factor as defined previously. Taking the Taylor expansion of Eq. (6) to estimate Δ*z*, we have

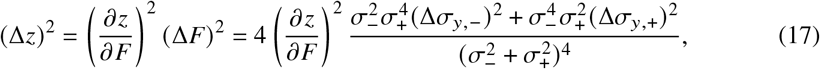

where 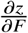 is the derivative of Eq. (6) computed numerically with our equation.

Considering the axial calibration curve shown in Fig. 2(c), the variation of the widths of the PSFs results in different axial precisions at different *z*-positions of the samples, giving us a minima at the center of the axial calibration curve when *z* = 0 nm. Here, we have the highest axial precision of 18.1 nm. Within the depth range of ±247.5 nm, the axial precision is sub-50 nm, which is an acceptable range of the biplane-based system (both 3D DWP- and 3D SDDWP-sSMLM) as highlighted in gray in Fig. 4(a). From Eq. (16), an increase in the photon count improves the estimate of the width of the PSF and the axial precision according to Eq. (17), as shown in Fig. 4(b).

**Fig. 4.**
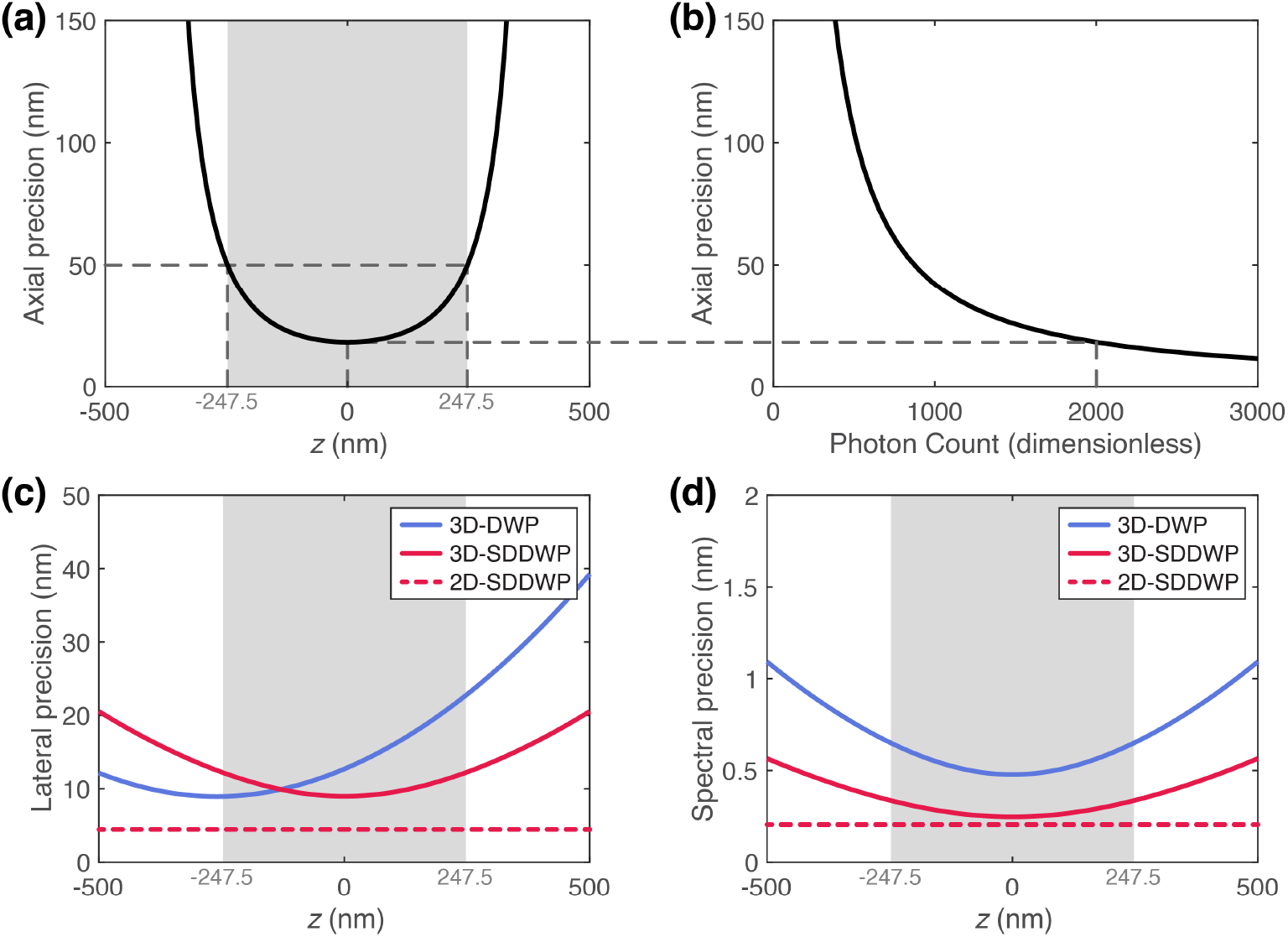
(a) Theoretical relationship between axial precision and depth position in 3D SDDWP-sSMLM; (b) Theoretical relationship between axial precision and photon count in 3D SDDWP-sSMLM; (c) Comparing the theoretical relationships between lateral precision and depth position in 2D SDDWP-, 3D SDDWP-, and 3D DWP-sSMLM; (d) Comparing the theoretical relationships between spectral precision and depth position in 2D SDDWP-, 3D SDDWP-, and 3D DWP-sSMLM.

#### 3.3.2. Lateral precision

The lateral precision in 3D DWP-sSMLM relies solely on the 0^th^ order image. As shown in Fig. 2(c), the minimum width of the PSF is observed at −250 nm on the axial calibration curve, which corresponds to the highest lateral precision since the precision is directly proportional to the width of the PSF according to Eq. (7). Consequently, the lateral precision of 3D DWP-sSMLM is optimal at 9.0 nm when *z* = −250 nm, and worse everywhere, as shown in Fig. 4(c).

In 3D SDDWP-sSMLM, lateral precision is determined by both the −1^st^ and +1^st^ order images. As shown in Fig. 4(c), the highest precision of 9.0 nm is at *z* = 0 nm. Overall, 3D SDDWP-sSMLM exhibits a 50% decrease in lateral precision compared to 2D SDDWP-sSMLM.

Within the depth range that the axial precision better than 50 nm, as highlighted in grey in Fig. 4(c), the average lateral precision of 3D DWP-sSMLM is 13.7 nm. In comparison, 3D SDDWP-system shows improved performance at 10.1 nm, demonstrating a 27% improvement.

#### 3.3.3. Spectral precision

In DWP- and SDDWP-SMLM, the spectral information associated with each localization is derived from the *x*-positions in the images from both orders. When PSF size increases with defocusing in biplane imaging, the localization precision of the *x*-positions worsens, as indicated by Eq. (7). According to Eq. (9), the spectral precision degrades further with increasing defocus, similar to that of lateral precision in 3D SDDWP-sSMLM. Fig. 4(d) shows the highest spectral precisions of 0.5 nm and 0.2 nm for 3D DWP- and 3D SDDWP-sSMLM, respectively, at *z* = 0 nm. The highest spectral precision in 3D SDDWP-sSMLM approaches the performance of 2D SDDWP-sSMLM. Within the depth range of −247.5 nm to 247.5 nm, where the axial precision is higher than 50 nm (highlighted grey in Fig. 4(d)), the average spectral precisions are 0.5 nm and 0.3 nm for 3D DWP- and 3D SDDWP-sSMLM, respectively; hence, 3D SDDWP-sSMLM exhibits a 48% improvement over 3D DWP-sSMLM.

### 3.4. Cellular imaging with 3D SDDWP-sSMLM

To demonstrate the feasibility of 3D SDDWP-sSMLM for cellular imaging, we imaged then processed HeLa cells with peroxisomes, microtubules, and mitochondria labeled by DY-634, AF647, and CF660C dyes, respectively, following the methods described in Sections 2.3.2 and 2.4.4. Figure 5(a) shows a depth-coded reconstruction of HeLa cells, and Fig. 5(b) shows a color-coded reconstruction of the different cellular structures. We observe that the mitochondria span over the entire *z*-depth in Fig. 5(a), which can also be visualized in Fig. 5(c), where the heatmap of the spectral peaks is plotted against the *z*-depth. Similarly, the microtubule structures identified in Fig. 5(b) are clearly visible in Fig. 5(c) to be concentrated towards lower *z*-depths, consistent with the observations in Fig. 5(a). Since the peroxisomes are not dense within the cells, they do not show up significantly in Fig. 5(c), but we see them correctly localized and visualized in Fig. 5(b). In Fig. 5(c), we see a distinct separation of AF647 and CF660C, despite the two having relatively highly overlapped spectra (Fig. S4(a)). DY-634 has a similar degree of spectral overlap with AF647, but due to the LPF in the emission path (Fig. 1(a)), some of the shorter wavelengths of the emission are being cut off, resulting in a higher emission peak (Fig. S4(b)), and leading to poorer spectral separation. We see low crosstalk between the channels in Fig. S4(c-e). In Fig. S4(f), we show a spectral maximum-encoded image of the same image, validating that the magnification corrections applied to minimize the bias to the calculated spectral peak independent of lateral position.

**Fig. 5.**
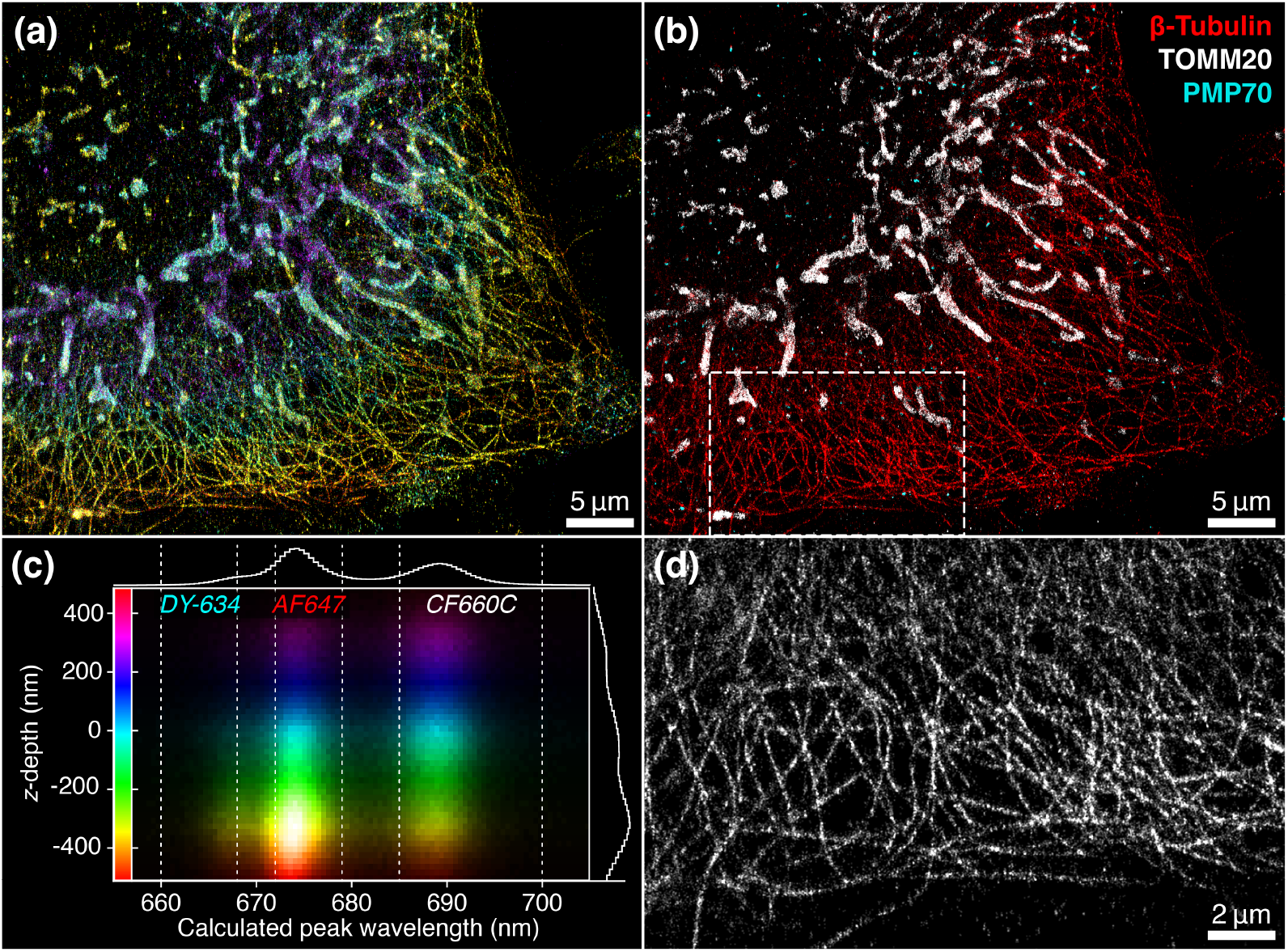
(a) Depth-coded reconstruction of HeLa cells with peroxisomes, microtubules, and mitochondria labeled with DY-634, AF647, and CF660C, respectively; (b) Color-coded rendering of HeLa cells with peroxisomes (cyan), microtubules (red), and mitochondria (white) labeled; (c) Heatmap of the calculated peak wavelengths of the image against *z*-depth. The histogram to the right represents the axial counts of the image in (a), while the histogram on the top represents the spectral profile of the image before reconstruction. (d) Magnified view of the microtubule structure from the highlighted area in (b).

Using three *z*-offsets to reconstruct an image adds to the overall depth that we can use for visualization, but we note two features from the *z*-depth histogram on the right of Fig. 5(c) that there are three peaks in the histogram, and the peaks are less prominent as we move up from *z* = −450 nm to *z* = 450 nm. First, the three peaks correspond to the focus positions of the three images respectively. From Fig. 4(c), we see that the precision is best at the *z* = 0 nm position of the image, which corresponds to the point where there is the highest signal-to-noise due to the least total defocus in both orders. With lower signal-to-noise ratios, fewer localizations can be identified; thus, the number of localizations decreases away from *z* = 0 nm. Next, the number of localizations taper off towards the end of the imaging sequence due to photobleaching over time. This causes the number of fluorophores available for imaging to decrease, resulting in fewer localizations over time.

Lastly, Fig. 5(d) provides a magnified view of the microtubule network, further demonstrating the system’s high spatial resolution. Notably, the absence of significant artifacts between microtubules in both lateral directions indicates good lateral precision. Overall, the results demonstrate 3D SDDWP-sSMLM’s capability to capture complex biological structures with high spatial and spectral precision, making it a powerful candidate for studying biological interactions at the nanoscale.

## 4. Conclusion

In this work, we have demonstrated the feasibility of 3D SDDWP-sSMLM for biological imaging. By incorporating two DWPs to disperse photons to the −1^st^ and +1^st^ orders symmetrically, we have significantly improved spatial and spectral precision compared to DWP-sSMLM. With a photon budget of 2,000, we have achieved a lateral precision of 10.1 nm, an axial precision of 18.1 nm, and a spectral precision of 0.3 nm. These results represent a 28% improvement in spatial precision, a 48% improvement in spectral precision, and a comparable axial precision compared with DWP-sSMLM. We have also demonstrated the capability to image complex biological structures with high spatial and spectral precision.

## Acknowledgments

Wei-Hong Yeo thanks the Christina Enroth-Cugell and David Cugell Fellowship for Visual Neuroscience and Biomedical Engineering at Northwestern University for their support.

## Data availability

Data underlying the results presented in this paper are not publicly available at this time but may be obtained from the authors upon reasonable request.

## Supplemental document

See Supplement 1 for supporting content.

