## Supplement 1 for "Maximizing photon utilization in spectroscopic single-molecule localization microscopy using symmetrically dispersed dual-wedge prisms"

### SPECTRAL CALIBRATION

Figure S2 shows an analysis of the spectral calibration parameters of dual-wedge prism (DWP)- and symmetrically dispersed DWP (SDDWP)-sSMLM. First, we localize our nanohole array (as portrayed in Fig. S2(a)) at different wavelengths, and then compute the mean $x$- and $y$-positions of the nanoholes. This gives us the mean position of the nanohole array at different wavelengths for both DWP- and SDDWP-sSMLM, and the resulting graph shown in Figs. S2(a) and (b), respectively.

To correct for the non-linearity due to the DWP, we use the invertible rational function provided in [1]. This gives us a linear relation between the mean localization of the nanohole array and the function $f(\lambda)$, where $f(\lambda)$ is the function that maps the wavelength to a corresponding scaled $x$-position of the nanohole array. We plot the mean localization of the nanohole array against $f(\lambda)$ for the $-1^{st}$ and $+1^{st}$ order images for both DWP- and SDDWP-sSMLM are shown in Figs. S2(c-f). From Fig. S2(c), we see that for the $-1^{st}$ order image of DWP-sSMLM, slope of the linear fit is close to zero, while in Fig. S2(d) for SDDWP-sSMLM, the slope is close to the value in the $+1^{st}$ order images of both DWP- and SDDWP-sSMLM. This is because there is a DWP at the beampath of the $-1^{st}$ order image of SDDWP-sSMLM, while there is no DWP at the beampath of the $-1^{st}$ order image of DWP-sSMLM. We also observe that the gradient of the two orders in SDDWP-sSMLM is not exactly equal, but this is due to some lateral shifts when performing the spectral calibration procedure. To account for lateral shifts in the image, we instead fit our equations to the x-shifts of the $-1^{st}$ and $+1^{st}$ order images.

The $x$-shifts of the $-1^{st}$ and $+1^{st}$ order images against $f(\lambda)$ for both DWP- and SDDWP-sSMLM are shown in Figs. S2(g) and (h), respectively. The slope of the linear fit, $m$, is given to be $-37.6$ and $-72.7$ for the DWP- and SDDWP-sSMLM systems, respectively, and is directly related to the spectral dispersion of the system. Thus, SDDWP-sSMLM provides double the spectral dispersion of DWP-sSMLM, due to the combined effects of both DWPs in SDDWP-sSMLM.

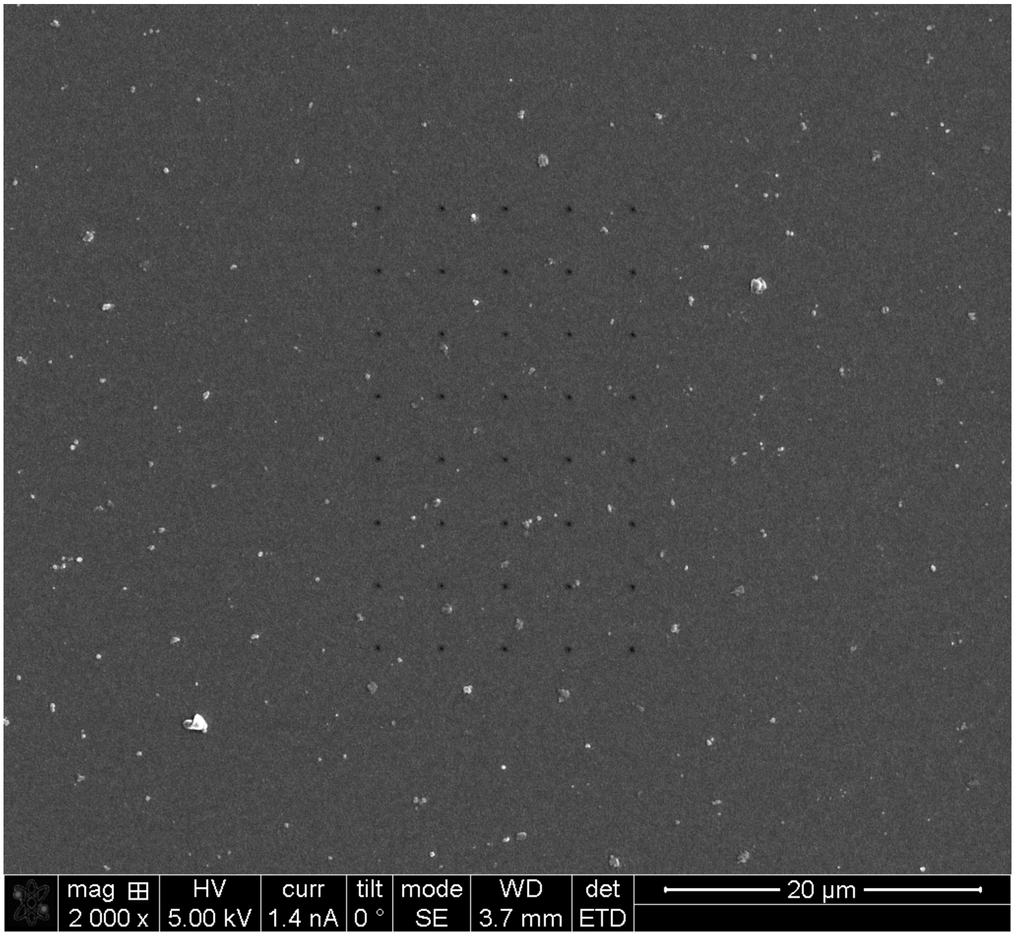


**Fig. S1.** SEM image of the nanohole array used for spectral calibration.

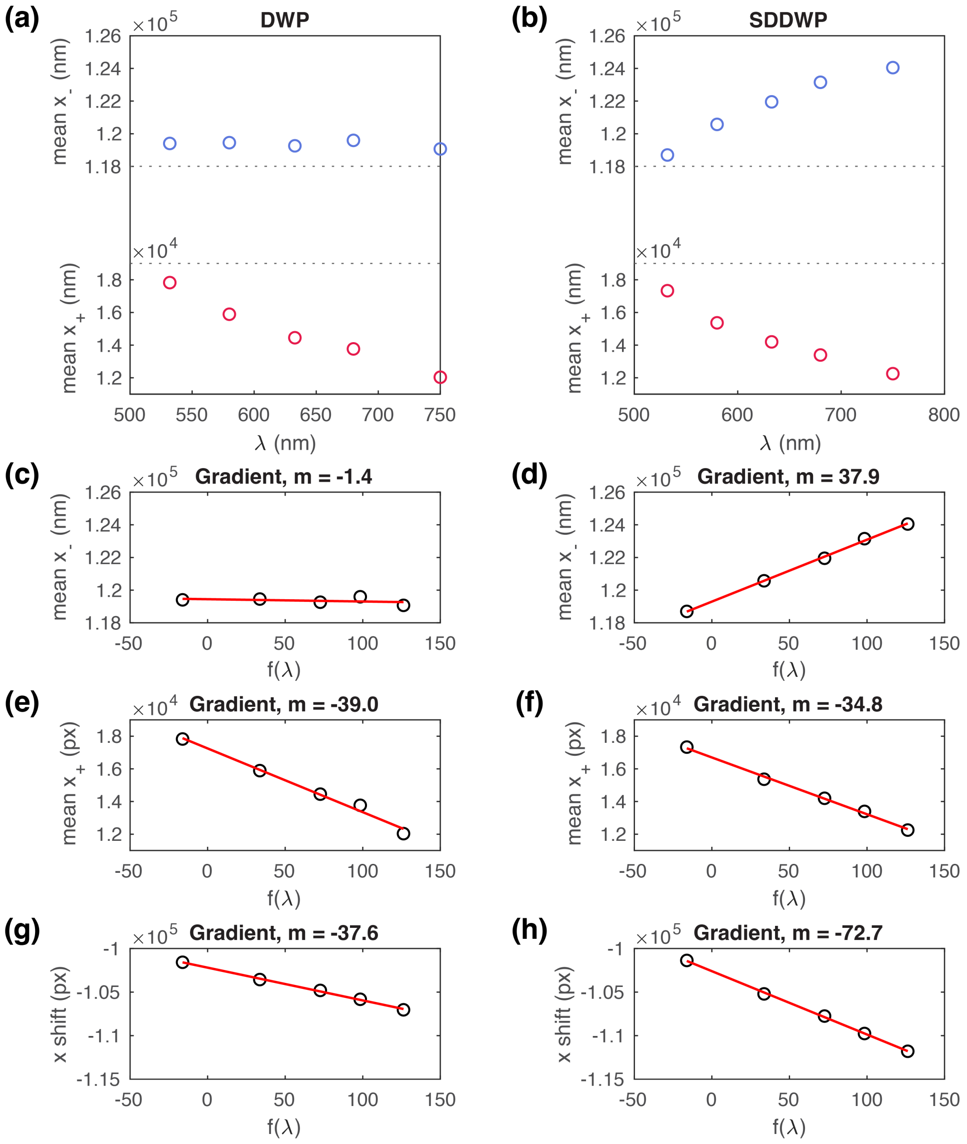


**Fig. S2.** Spectral calibration of the dual-wedge prism system. (a-b) Mean localizations of the nanohole array at different wavelengths for (a) DWP- and (b) SDDWP-sSMLM. (c-f) Plot of the mean localization of the nanohole array against $f(\lambda)$ for the $-1^{st}$ and $+1^{st}$ order images, for both (c,e) DWP- and (d,f) SDDWP-sSMLM. The solid lines are linear fits to the data. The slopes of the linear fits are used to calculate the spectral calibration coefficients. (g-h) Plots of the $x$-shifts of the $-1^{st}$ and $+1^{st}$ order images against $f(\lambda)$ for (g) DWP- and (h) SDDWP-sSMLM.

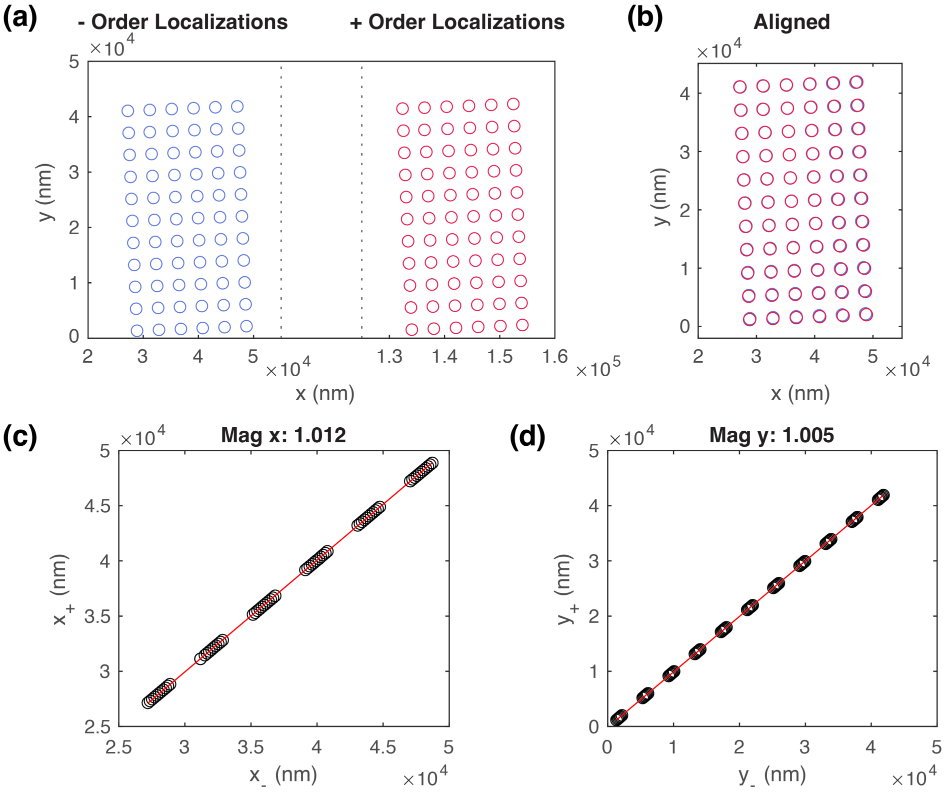


**Fig. S3.** Magnification calibration of the dual-wedge prism system. (a) The raw localizations of image from the SDDWP-sSMLM system. (b) Aligned localizations by subtracting the difference in the mean locations between the $-1^{\mathrm{st}}$ and $+1^{\mathrm{st}}$ order images. (c) A plot of the $x$-positions of the $-1^{\mathrm{st}}$ against the $+1^{\mathrm{st}}$ order images. (d) Plot of the $y$-positions of the $-1^{\mathrm{st}}$ against the $+1^{\mathrm{st}}$ order images. The solid line is a linear fit to the data.

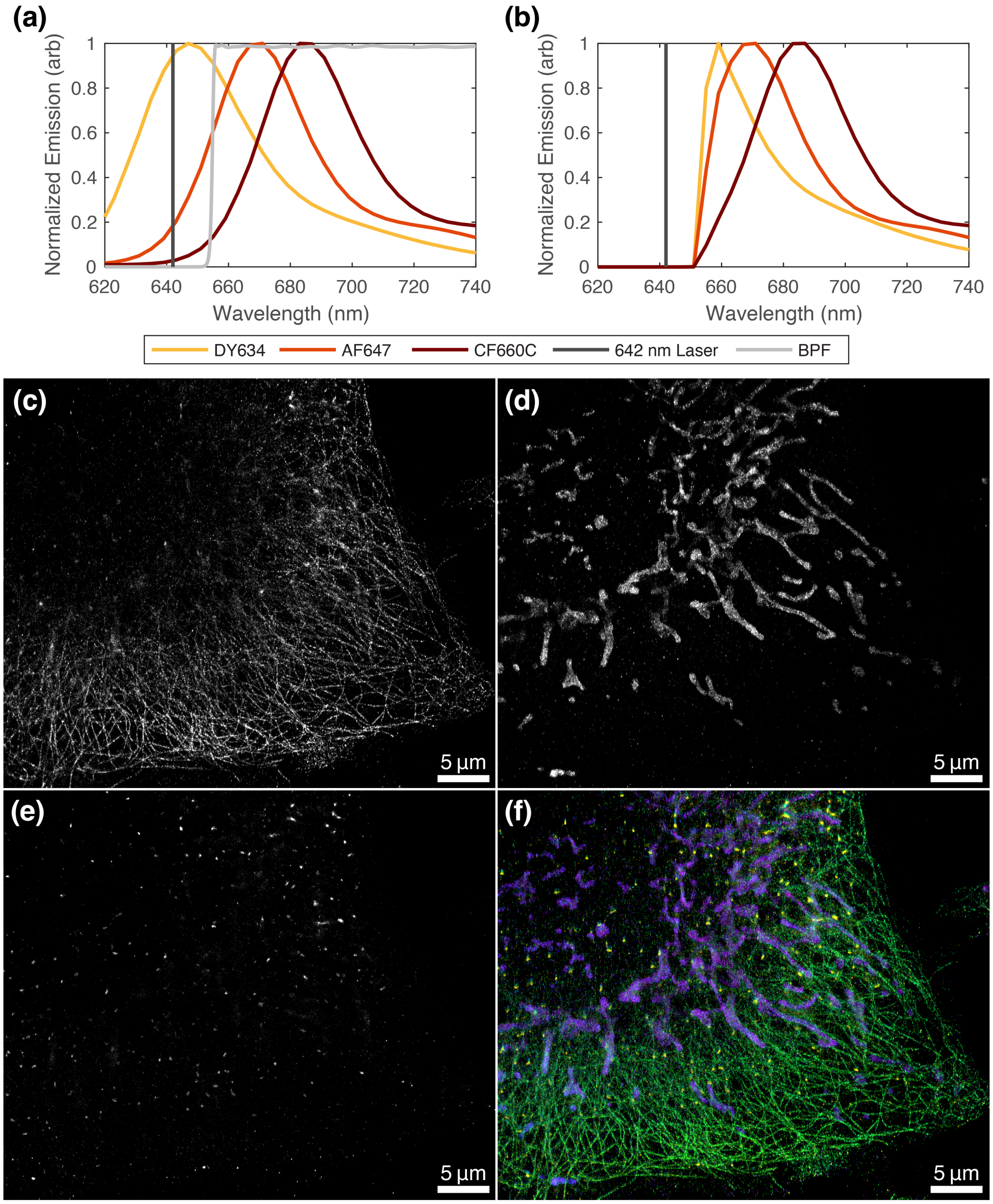


**Fig. S4.** (a) Plots of DY-634, AF647, and CF660C dyes extracted from bulk data measured on Nanodrop Spectrofluorometer. The grey line shows the LPF transmission data taken from Semrock. (b) The spectral peaks of the dyes after normalization with the transmission efficiency of the LPF. Individual channel images of the (c) microtubules, (d) mitochondria, and (e) peroxisomes. (f) We show a color-coded image of the calculated spectral peak of the cell image. Visually, there is clear separation of the different features, suggesting good spectral resolution with excellent field of view.
